# Adaptive Coding Explains the Paradoxical Relationship between hippocampus size and memory function in Alzheimer’s Disease, Autism Spectrum Disorder, and PTSD

**DOI:** 10.1101/2023.11.22.568352

**Authors:** Andrea Stocco, Holly S. Hake, Bridget Leonard, Xinyue Li

## Abstract

The relationship between hippocampal volume and memory function has produced mixed results in neuroscience research, which suggests an additional influencing mechanism. To explore the role of an experience-dependent encoding mechanism, we developed an autoencoder model of the cortex-hippocampus loop and examined how its memory representation is affected by a penalty that prioritizes sparseness. We trained our model with the Fashion MNIST database and a loss function to modify synapses via backpropagation of mean squared recall error. The model exhibited experience-dependent efficient encoding, representing frequently-repeated objects with fewer neurons and smaller loss penalties; representations of similar sizes were found for objects repeated equally. Our findings clarify perplexing results from neurodevelopmental studies by linking increased hippocampal size and memory impairments in autism spectrum disorder (ASD) to decreased sparseness, and explaining dementia symptoms of forgetting with varied neuronal integrity. Our findings propose a novel model that connects observed relationships between hippocampal size and memory to the environmental demands and the underlying biological constraints, contributing to the development of a larger theory on experience-dependent encoding and storage and its failure.

**Author Summary:** The hippocampus is the brain region where memories are initially formed. A larger hippocampus seems to be associated with better memory performance, but this is not a static relationship; studies suggest that its size might grows with demand (e.g., the need to memorize more information) and that, in certain conditions, larger or smaller volumes of the hippocampus are not associated with better or worse memory. We demonstrate that it is possible to make sense of these findings by assuming that the hippocampus balances the need to correctly remember with the need to minimize resources used to store it, and that the hippocampal size might change as a function of memory demands (the characteristics of the information to memorize) and the underlying biology (e.g., the presence of hippocampal cell damage, or a reduction in hippocampal GABA receptors). We test this hypothesis by building a neural network model; the model correctly predicts existing, puzzling findings in the literature, and can even capture subtle interactions between different phenomena, such as predicting that individuals with Autism Spectrum Disorder would be more susceptible to memory loss in dementia.

## Introduction

The hippocampus is a region of the medial temporal lobe that is critical for long-term memory storage and retrieval. The size of the hippocampus can vary significantly between individuals and these variations in size have been associated with corresponding differences in memory function (Pohlack et al. 2014; Hardcastle et al. 2020; Botdorf, Canada, and Riggins 2022). This relationship between hippocampal size and memory function, however, is complex and not always straightforward [1]. There is evidence that greater hippocampal volume is associated with better memory function; for example, greater hippocampal volume is associated with longer-lasting spatial memory [2], and better performance in spatial memory tasks [3], and superior performance in a variety of verbal memory tasks [4]. Conversely, reduced hippocampal size is often associated with significant impairments in long-term memory: this is the case of fronto-temporal degeneration and Alzheimer’s disease, in which neuronal loss results in markedly reduced hippocampal volume, and the degree of volume loss positively correlates with the severity of amnestic symptoms [5,6].

Nevertheless, the relationship between hippocampal size and memory performance is also, at least partially, plastic and mediated by experience. A notable and oft-reported case is the fact that London taxi-cab drivers have a larger hippocampus than the normal population. This has been speculated to be connected to the amount of spatial information that cab drivers need to memorize (“The Knowledge”) to exercise their profession [7]. In fact, a follow-up study revealed that changes in hippocampal size follow, and do not precede, the amount of studying necessary to pass the test [8]. Similarly, changes in hippocampal size correlate with an individual’s years of education [9].

An intuitively appealing explanation for these effects might be that the hippocampus grows with the environmental demands—that is, the amount of data it needs to, or can, store. Thus, the process of storing more information results in the growth of the hippocampus, and a reduction in hippocampal size results in loss of memory. At the same time, the size of the hippocampus constrains the amount of knowledge that can be stored. Thus, pathologically reduced hippocampal volume implies less room to store memories, and a larger hippocampus might confer greater memory capacity.

This simple and intuitive explanation, however, is complicated by a number of other findings. Reductions in hippocampal size are common in a variety of mental disorders, including posttraumatic stress disorder (PTSD: [10]) and, to a lesser extent, anxiety disorders. when executive functions are accounted for, reductions in HPC size is not accompanied by changes in long term memory functions [11]. Conversely, larger hippocampal volume has been observed in autism spectrum disorder (ASD), where a corresponding increase in memory function was not observed; in fact, individuals with ASD exhibit slight *deficits* in episodic memory [12,13]. Furthermore, a recent epidemiological study found that the prevalence of amnestic forms of dementia in ASD is up to four times higher than the neurotypical average [14], despite the fact that greater hippocampal size could have represented a buffering factor against neuronal loss. Thus, while it has been shown that experience drives changes in hippocampal size, changes have also been observed in clinical conditions without corresponding changes in memory—or with changes in the opposite direction.

At least three possible explanations can be proposed to reconcile these findings. The first, and perhaps the most mundane, is that changes in hippocampal size might not always reflect underlying changes in the number of hippocampal cells or synapses. Virtually all of the studies reported here assess hippocampal size through anatomical MRI, and the sheer volume of a region in an MRI scan can be affected by a variety of other factors. For instance, greater water density in tissue also leads to greater volumes but is not associated with a larger number of cells or synapses [15].

A second explanation is that two or more biological mechanisms might be at play simultaneously. Experience-dependent growth following intense memory training and dementia-related loss of memory function are both linked to the number of cells and synapses. However, changes in other clinical domains, such as those seen in PTSD and anxiety, might stem from distinct mechanisms. For example, prolonged stress exposure causes neuronal death through the accumulation of cortisol, resulting in volume loss. Thus, it is possible that the volume loss in PTSD and anxiety are due to stress-induced cortisol-related pruning [16,17], which does not play a role in dementia or ASD.

The third and last explanation is that these distinct phenomena are indeed connected by experience-dependent efficient allocation of hippocampal cells and synapses to varying memory demands, but that this relationship is complex and non-linear.

In this paper, we put forward a neurocomputational framework that accounts for this third hypothesis. According to this framework, the need to store and retrieve memories demands an efficient allocation of neural resources; it is this need for efficiency that mediates between neuroanatomical constraints (such as loss of neurons) and environmental demands (such as greater need to memorize information, or the presence of persistent and intrusive memories). In turn, the principles underlying the efficient allocation of resources can be understood in terms of information theory.

The remainder of the paper is structured as follows. First, we provide a theoretical foundation for our framework in terms of the Free Energy Principle [18,19] and a review of previous attempts to model the relationship between memory function and hippocampal size. Next, we introduce a neural network model, based on an autoencoder architecture, that demonstrates how the hippocampus could adaptively allocate neurons in response to varying demands under realistic conditions. Finally, we speculate on the implications of this mechanism for two important memory-related phenomena: sleep and spontaneous brain activity.

### Relationship to Free Energy and Predictive Coding

The Free Energy Principle (FEP) is a general-purpose framework to interpret brain activity. According to the FEP, the brain’s main goal is to minimize the *surprisal* of future environmental stimuli; formally, the surprisal of a given stimulus *S* is computed as –log_2_ *P*(*S*), with *P*(*S*) being the probability of *S* occurring in the environment. In information theory, the surprisal is also known as the *information content* of *S*. The rationale behind the FEP is simple: by maximizing the predictability of future stimuli, an organism can both maximize the opportunities for survival and minimize the metabolic costs associated with responding to the environment. In this sense, the FEP can also be seen as a mathematical formulation of the idea of *predictive coding* [20,21], that is, the idea that feedforward networks can be trained by minimizing each layer’s error in predicting its inputs from its feeding layers.

The FEP can be interpreted as constraining the *representations* that are learned within a population of neurons (although the concept of representation is vague [22]). Assuming that neural representations are sparse (i.e., only a subset of neurons are used to represent any specific stimulus) and distributed (i.e., the same neuron could be used in multiple representations), this interpretation of the FEP entails that the number of neurons that are spent to represent a stimulus should be proportional to its surprisal –log_2_*P*(*S*), (i.e., novel or unexpected stimuli would have the largest neuronal representation). This is, in fact, a specific implementation of the idea of efficient coding [23].

In this paper, rather than considering the representation of an incoming stimulus *S*, we will consider the representational costs associated with storing its memory *m*. Assuming the brain’s episodic memory system follows the FEP, it is possible to estimate the number of neurons used to represent all memories *m*_1_, *m*_2_, *m*_3_ … *m_M_* that have been encoded. First, according to the FEP, the number of neurons used to represent each memory would be proportional to –log_2_ *P*(*m*), with *P*(*m*) being the *need probability* of *m*, that is, the probability that *m* would need to be retrieved at a given time [24]. Second, because the representations are distributed, we need to compute the probability that a given neuron belongs to a single representation rather than being shared by others. In turn, this probability should be proportional to *P*(*m*): the more probable a memory *m* is, the smaller its representation, and the more likely it is that its neurons can be uniquely used for *m*. Thus, the total number *N* of neurons that can be used to represent all memories is proportional to:

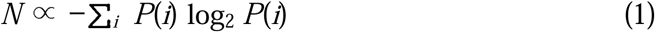

Note that the quantity –∑*_i_ P*(*i*) log *P*(*i*) is the definition of Shannon’s information entropy *H*, or information content, of all the memories [25]. *H* quantifies the average amount of surprisal or uncertainty involved in predicting the outcome of a random variable, or, in this case, in predicting the next stimulus based on the accumulated memories.

Equation 1 can be immediately used to explain experience-dependent changes in the hippocampus, such as those reported for cab drivers by Maguire and colleagues [8]. The total number of streets and landmarks in London, encoded in the Knowledge test, is approximately 75,000, and a taxi cab driver has to memorize them all to prepare for the test. Because all streets and landmarks are equally likely to be used in the test, the candidates need to practice all items equally, resulting in a roughly uniform distribution of the probabilities that each item would be needed. Uniform distributions are precisely the ones that maximize the entropy *H*. In contrast, a bus driver ony needs to memorize a small subset of these streets and landmarks; therefore, the distribution of their spatial memories will deviate significantly from uniform, with only a few streets and landmarks being retrieved and encoded very frequently (and thus having a high need probability) and some being scarcely used. In fact, it is not even necessary for a bus driver to know *all* of the 75,000 streets and landmarks; for simplicity, however, we will just assume that all such items have been encoded at least once, and thus have a small but non-zero need probability. Figure 1 compares a uniform distribution with three possible non-uniform distributions: linear, quadratic, and normal. It is easy to see that, no matter what, the entropy associated with the type of study carried out by taxi cab drivers, where each item is equally likely to be tested, is maximized compared to any other distribution of knowledge.

**Figure 1:**
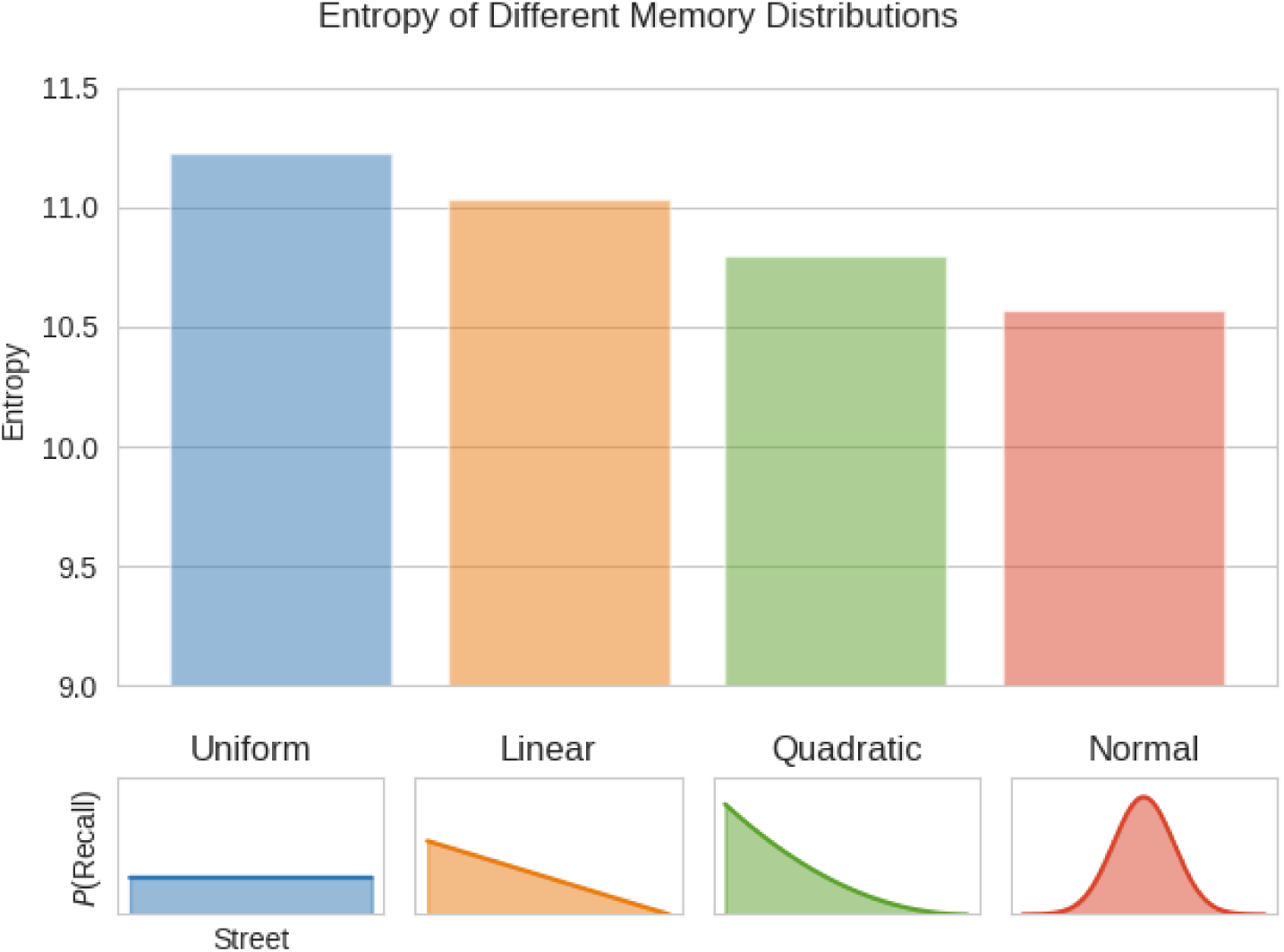
Entropy across four different possible distributions of the need probabilities for the 75,000 London streets. The Uniform distribution (left, blue) approximates the distribution of memories for cab drivers taking The Knowledge test. Possible distributions for the memories of bus drivers have (orange, green, and red) are associated with significantly less entropy.

### Previous Work

To the best of our knowledge, the first computational account of the relationship between memory demands and hippocampal size was put forward by Smith et al. [26]. The authors based their work on the framework originally proposed by Anderson and Schooler [27] and currently implemented in the ACT-R architecture [28]. In this framework, each memory is a collection of episodic traces, and each trace corresponds to a different encounter of the same information. The strength of each trace decays over time according to a power function with exponent *d*, and the memory’s total strength, or *activation*, is the log of the sum of its traces:

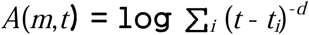

The probability *P*(*m*) of retrieving a memory can be computed as a function of its activation, relative to all other memories:

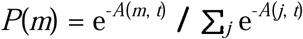

Smith et al. [26] proposed that the distribution of probabilities across all memories could be used to predict changes in hippocampal volume in PTSD. Specifically, the intrusive nature of traumatic memories leads to an accumulation of their traces, leading to increased activation and skewing the probability distribution of retrievable memories, making them less uniform. The higher the probability of retrieving intrusive memories, the lower the entropy of the model’s memory system, and, as a correlate, the lower the volume of the hippocampus.

However, the original model by Smith et al. [26] did not include any biological mechanisms for efficiently allocating neurons to different memory representations. It also does not solve the challenge of how the hippocampus could form an adaptive engram without prior knowledge of future activation patterns. Moreover, while the model accounts for changes in hippocampal volume specifically related to PTSD, it does not consider other experience-dependent changes, such as those associated with learning, nor does it address other findings— like the increased hippocampal volume observed in ASD or the link between reduced hippocampal size and memory decline in dementia. These limitations suggest a need for a more comprehensive model that can incorporate both pathological and experience-driven variations in hippocampal structure and function.

## An Autoencoder Model of the Hippocampus

This paper addresses these limitations by examining the behavior of a neural network model of the hippocampus and, specifically, testing whether (a) Efficient coding and efficient allocation of resources spontaneously emerges through the interaction of training the network and the network’s constraints; and (b) Whether the model can account for the diversity of findings relating hippocampal size and memory function.

### Architecture of the Model

The model was designed to capture the connectivity between cortical regions and the hippocampus so that both encoding and recall could be simulated. The connections between the cortex and the hippocampus form a recurrent loop. The exact set of synapses varies slightly across regions; as an example, this paper will consider the connectivity between the inferior temporal cortex and the hippocampus. This specific circuit is well understood and underlies memory for higher-level visual objects, which will also be used as experimental stimuli.

Projections from the inferior temporal cortex pass through the entorhinal cortex and the dentate gyrus and, through the mossy fibers, reaching area CA3 of the hippocampus; cells in this area bind converging multisensory inputs together, are are thus considered the initial locus of an engram [29,30].During recall, memories are then reactivated in the cortex [31,32] through a series of connections that originate in CA3 and progress through area CA1, the entorhinal cortex again, and finally return to the temporal cortex [33].

For convenience, this recurrent loop can be “unrolled” and visualized as a feedforward neural network with multiple layers. The first half of this network corresponds to the neural populations that respond to cortex-to-hippocampus projections during encoding, and the second half to the neural populations along the hippocampus-back-to-cortex projections during recall. In this design, the temporal cortex is both the input and the output of the network, and the hippocampus is the central bottleneck.

This network architecture is known as an autoencoder [34,35] and is used, in deep-learning applications, to learn a set of features that would efficiently compress the original input so that its output is minimally different from its input.

The model’s full architecture is shown in Figure 2. Its input is a 28×28 matrix that contains a grayscale visual representation of an object. This representation is then flattened to a layer of 784 neurons, which represents the object as encoded in the inferior temporal cortex. This representation is passed through a smaller layer of 512 neurons, representing the entorhinal cortex, and an even smaller one of 384 neurons, representing the dentate gyrus. It finally reaches a layer of 256 neurons that hold a compression representation of the original content and stands for the hippocampus’ CA3 field. Thus, the decreasing size of each layer mimics the relative sizes of the corresponding biological structure.The output of the hippocampus is then passed through a mirror series of layers representing CA1, the entorhinal cortex, and the visual cortex again (784 neurons), generating a reconstructed version of the original stimulus. All of the neurons in the model use a Rectified Linear Unit (ReLU) activation function, with the exception of the very last layer, which uses a sigmoid activation function to ensure that all of the predicted grayscale intensities fall between 0 and 1.

**Figure 2:**
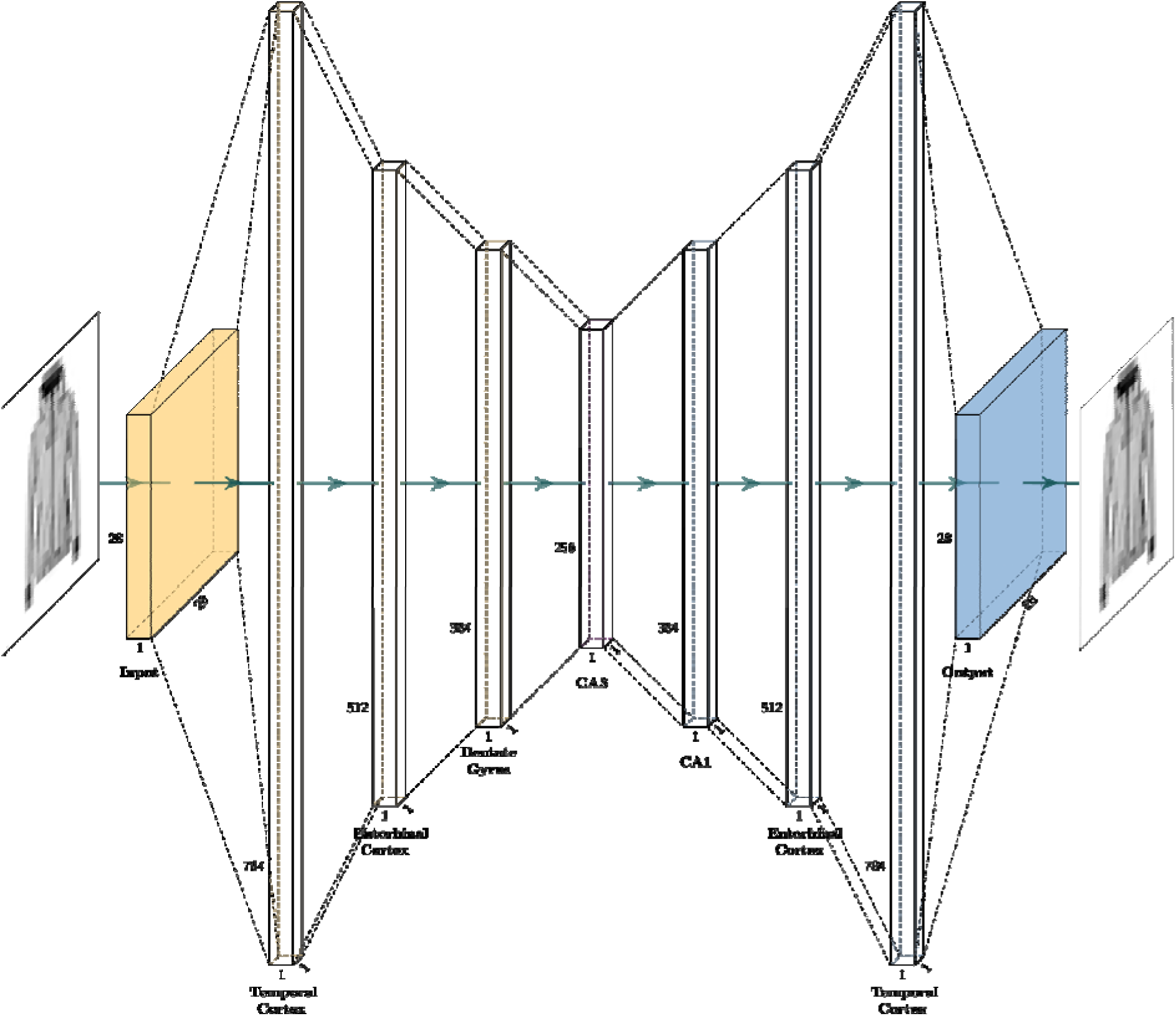
Architecture of the autoencoder neural network model. The first layers of the model (in yellow) represent the encoder, while the last layers of the model (in blue) represent the decoder. Area CA3 of the hippocampus is depicted in purple.

### Design Choices and Their Relationship to the Existing Literature

The proposed model differs significantly from the majority of computational models that have been proposed for the hippocampus. Most of these models [31,36,37] treat the hippocampus, and especially region CA3 as an *autoassociator*, a network capable of content addressable storage. This hypothesis is supported by the neuroarchitecture of CA3, with most neurons forming excitatory recurrent synapses with other CA3 neurons [38]. Few models, however, have explored the nature of connections between CA3 and the cortex (see O’Reilly and Treves for an exception). These connections are, in fact, essential to better understand the interplay of encoding and recall. Furthermore, an important limitation of autoassociative models of the hippocampus is that their representations do not change over time and are, in general, unaffected by the frequency of the presentation of each stimulus. Thus, they are poorly suited for capturing the experience-dependent phenomena that are the focus of this study.

In contrast, in this model, the hippocampus and the surrounding areas form a hierarchical system that compresses information during encoding and fills it in during retrieval. To a large extent, the application of autoencoders in this model can be understood as mirroring the function of episodic memory and, by extension, the role of the hippocampus.

This interpretation is more closely aligned with the CRISP theory of the hippocampus [39]. Unlike the CRISP framework, however, our model does not include recurrent temporal dynamics; each encoding and retrieval act is performed in a single pass.

As mentioned above, most of the neurons are rectified linear units. ReLUs are the standard in modern deep-learning neural networks due to the computational advantages they offer, which often result in significant time savings in the training of the models. In this model, however, the choice of using ReLUs was made to ensure that we could unambiguously count neurons involved in representing a specific object—in this case, neurons whose output is greater than 0. Note that, by using ReLUs, we are also stacking the cards against our prediction, since the use of ReLUs provides an intrinsic, rather than data-driven, mechanism for sparsification.

Thus designed, this model was used in five sets of simulations, each of which addresses a different facet of the relationship between hippocampal size, memory representation, and memory function. In each simulation, the model was trained on a selection of objects from the Fashion MNIST database [40], a collection of 70,000 28×28 black-and-white images of 10 categories of clothing items (e.g., boots, pants, bags, shirts, …). This particular dataset was chosen because it is made of visual representations of realistic inanimate objects, such as those that would activate the inferior temporal cortex, at a relatively low stimulus resolution, thus reducing the computational load of the simulations. In every simulated run, the model was trained with a subset of images that were randomly selected. Each selection included 1,111 images, which were repeated with varying frequency across simulations (see below in *Simulation 1*) to always form a training set of 4,000 stimuli.

In all simulations, The model was trained on all items in the training set for five consecutive epochs. During training, stochastic gradient descent was applied using a combined loss function *L* that included two terms:

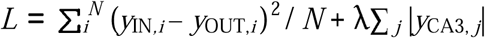

In which the notation *y*_L*,n*_ refers to the activation *y* of the *n*-th neuron in layer *L*. The first term is the *accuracy cost* of the network’s recall function, that is, the mean squared difference between the activations of each input neuron *y*_IN*,i*_^]^ and the activation of each corresponding output neuron *y*_OUT,*i*_. The second term is the *resource cost* and is the sum of the activations of each neuron in CA3; this term was added to increase the need for sparseness in the hippocampal representations. The hyperparameter λ regulates the weight of the penalty and was set to 0.00001 throughout these simulations. This value is somewhat arbitrary; however, preliminary simulations showed that the results did not qualitatively change for different values of lambda, as long as its value was below a critical threshold of 0.0001, above which the penalty became too severe.

## Results

### Simulation 1: Emergence of Efficient Coding

In the first simulation, the set of 1,111 objects was used to create a 4,000-item training set in which different objects were repeated with different frequencies. Specifically, 1,000 objects occurred only once, 100 objects occurred 10 times, 10 objects occurred 100 times, and a single object occurred 1,000 times. This simulation was run 50 times. If the model is learning a form of efficient coding, the internal hippocampal representation of an object should depend on its frequency in the training set, and, therefore, objects that are repeated the most should have representations with fewer neurons and smaller L1 penalties than objects that are repeated the least.

Figure 3 illustrates the results of one such simulation. The top row shows four example objects from one specific simulation, chosen from the sets of stimuli repeated 1, 10, 100, or 1,000 times, respectively. The middle row represents the corresponding responses of the simulated CA3 layer, with the activations of its 256 neurons arranged in a 16×16 grid. The dependent variables for hippocampus sparseness (L1 penalty and number of neurons) are also reported. Finally, the bottom row depicts the recalled memory. Note that, although recall is generally accurate, the precision and the level of details of the recalled memory might vary significantly across stimuli; with recall being exact for some of them (e.g., the tennis shoe) and imprecise for others (e.g., the *t*-shirt).

**Figure 3:**
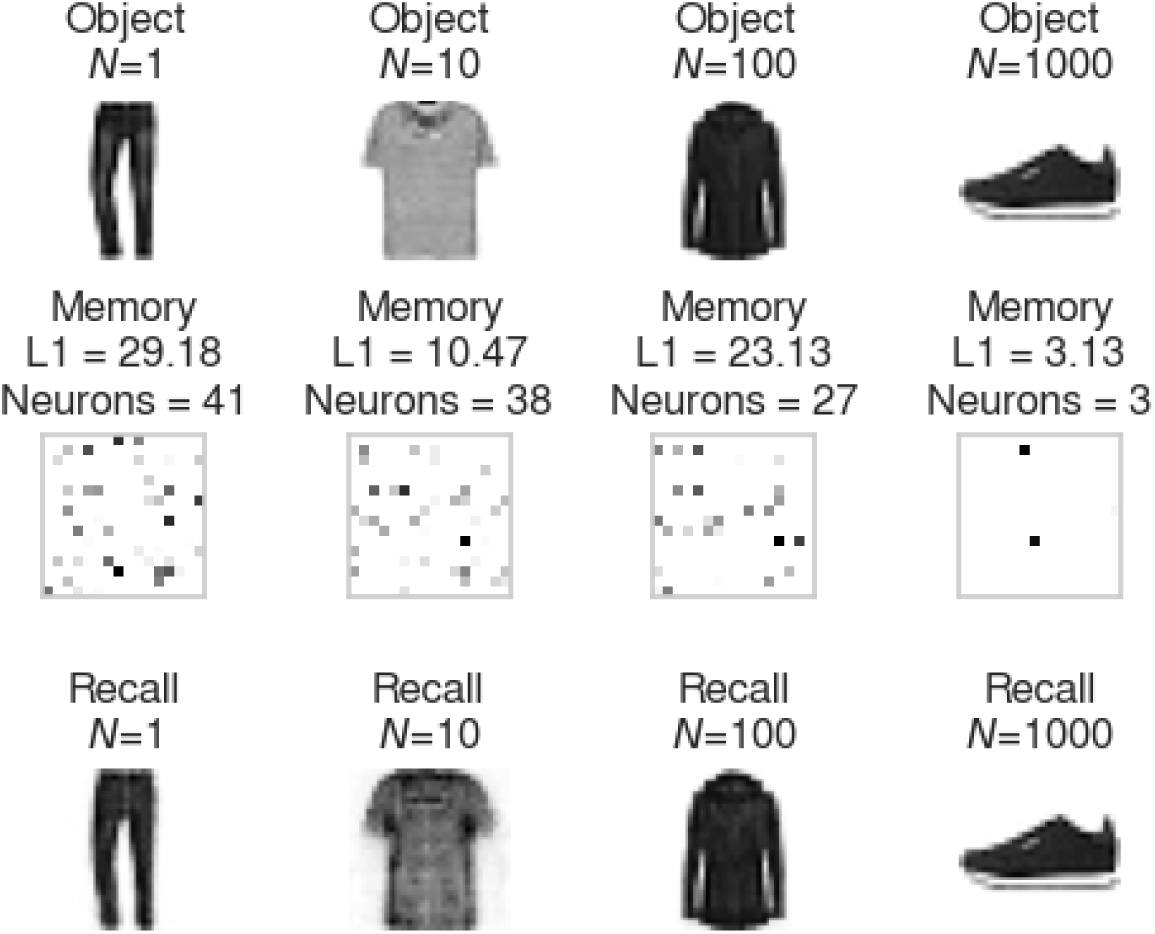
(Top) Four example stimuli that were repeated 1, 10, 100, or 1,000 times in the training set. (Middle) Corresponding CA3 representations of the stimuli; (Bottom) Recalled stimuli reconstructed by the decoder from the CA3 representations.

Although representative, Figure 3 only depicts four example objects from a single run. A complete overview of all the simulations is instead reported in Figure 4, where the mean L1 penalty and the mean number of neurons are reported as a function of the object frequency.

**Figure 4:**
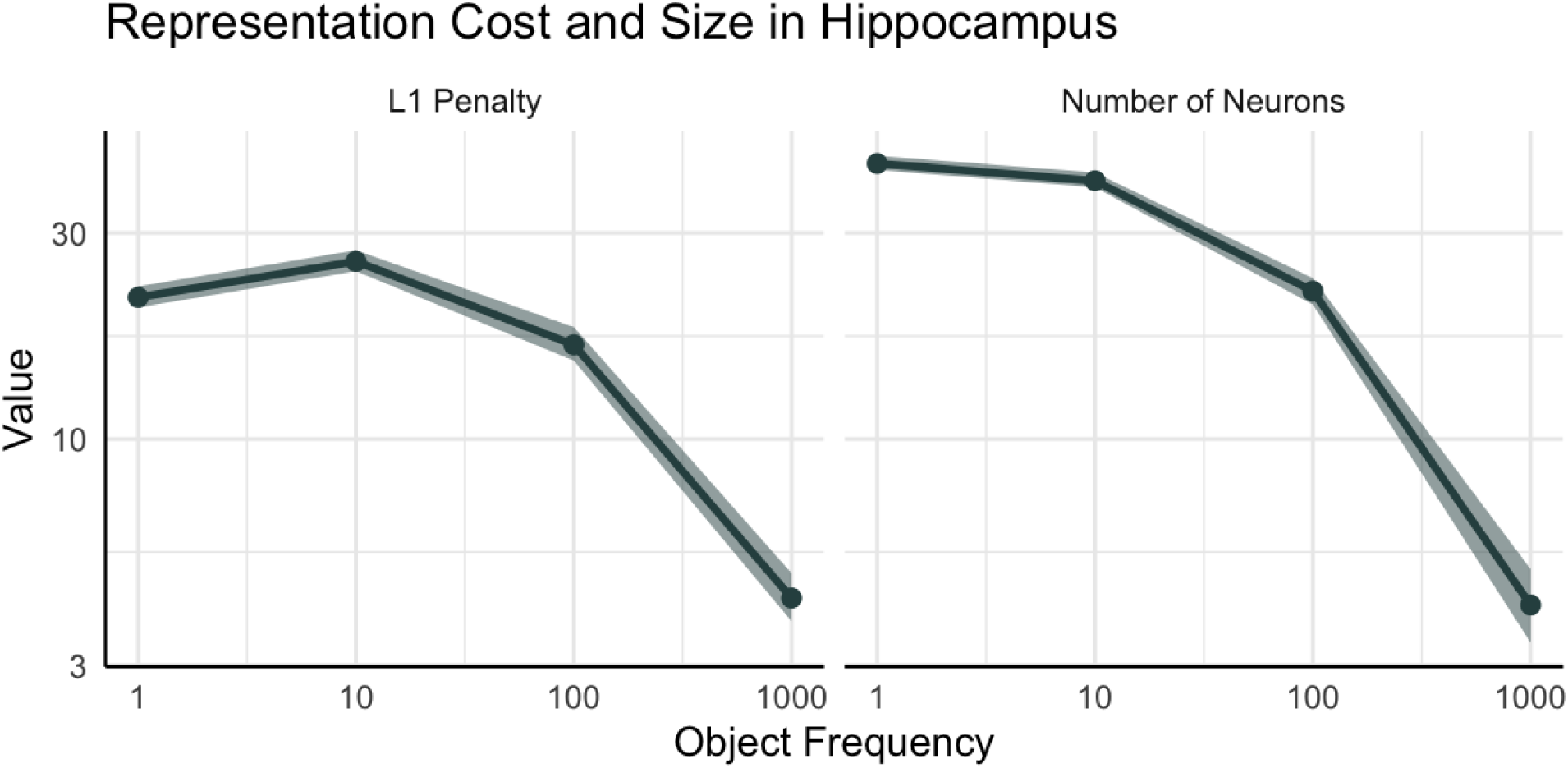
L1 penalty and number of neurons as a function of the frequency of the presented object. Lines represent means, ribbons represent 99% confidence intervals.

The effects of an object frequency are apparent for both measures. A statistical analysis was performed by fitting a linear regression model over the results of all 50 simulations, using object frequency as the independent variable and either L1 penalty or number of neurons as the dependent variable. The effect of frequency was significant for both outcomes (L1 penalty : β = –0.02, *p* <.0001; Number of neurons: β = –0.02, *p* <.0001).

#### Relationship to Information Theory

But how closely does the reduction in the CA3 representation match the predictions of information theory? According to Huffmann [23] efficient codes are such that the length of a code for an object *x* matches its information content or surprisal, that is, –log_2_ *P*(*x*). The value of *P*(*x*) can be calculated from the number of occurrences of stimulus *x* in the training set. Figure 5 compares the relationship between the number of neurons used to encode an object in the CA3 layer and its corresponding information content. As the figure shows, the number of neurons closely mirrors (*r* = .97, *p* =0.03) the information content, a hallmark of efficient coding.

**Figure 5:**
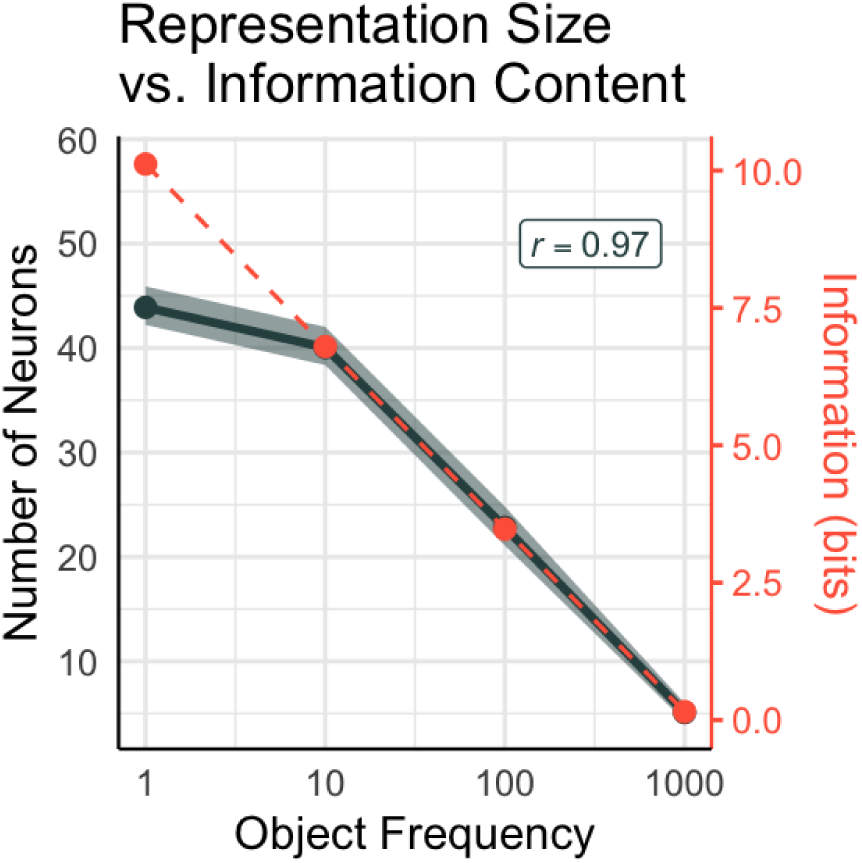
Relationship between the number of neurons that encode a stimulus (charcoal line and ribbon, representing mean +/– 99 confidence interval) and its information content (red line).

### Simulation 2: Object Frequency Drives Efficient Coding

Although the previous results provide convincing evidence that different object frequencies result in representations of different sizes, they do not provide causal evidence for the direction in which the process works. Hypothetically, the previous results could be explained by the model allocating longer codes to the less frequent memories, rather than shorter ones to the most frequent ones. To eliminate this possibility, a second set of simulations were run in which the model was trained with a dataset of identical size (4,000 items) and containing the same 1,111 objects, but in which each object was repeated an approximately equal number of times, either 3 or 4. In these simulations, objects maintained the original frequency label as in the previous one; for example, the 10 objects that were originally repeated 100 times were still classified as in the “100 repetitions” group, even if they were repeated only 3 or 4 times each.

Figure 6 depicts the results of these simulations (in red), plotted against the results of the previous simulations (in charcoal). Note that, when all objects appear at approximately the same frequency, both the L1 penalty and the number of neurons remain stable and at the same level of the least frequent memories in Simulation 1. (Note that, in this case the training set with equal frequencies, the values on the *x*-axis do not actually reflect the frequencies on the previous simulations’ training set). Regression models confirmed this, finding virtually no relationship between the original object frequency and the resulting L1 penalty (β < 0.001, *p* > .78) or number of neurons (β < 0.001, *p* > .80). Furthermore, a series of *T*-tests revealed that both the L1 penalty and the mean number of neurons were significantly *lower* when objects were presented 100 or 1,000 times than when their frequencies were equal (*T* > 5.06, *p* < 0.001, Bonferroni-corrected; marked by “*” in Figure 6) but not significantly different at frequencies of 1 or 10 repetitions (*T* < 2.5, *p* > 0.05, Bonferroni corrected). Thus, our model demonstrates that increased presentation frequency does result in sparser hippocampal representations of the corresponding object.

**Figure 6:**
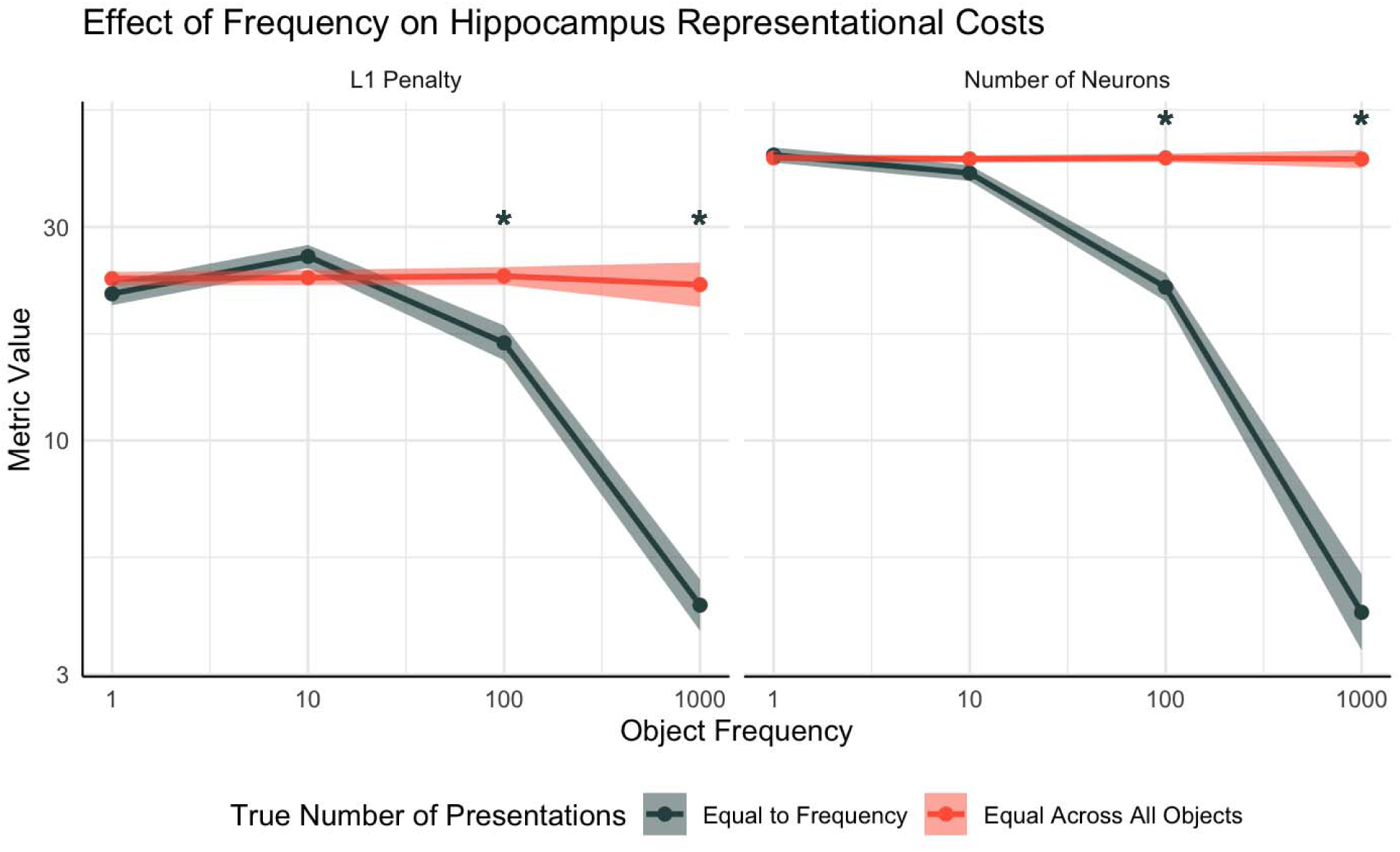
L1 penalty and number of neurons as a function of the frequency of the presented object. Lines represent means, ribbons represent 99% confidence intervals. Simulation 1 had varied frequencies (charcoal) and Simulation 2 had equal frequencies (red line). Asterisks denote object frequencies for which a significant difference (*p* < 0.05, Bonferroni corrected) was observed between the two simulations.

#### Experience-dependent Changes in Hippocampal Size

These results suggest a possible computational mechanism for changes in hippocampal size that are dependent on experience, such as those observed in London taxi cab drivers compared to bus drivers [8]: as noted by Smith et al. [26], unlike bus drivers, taxi cab drivers must rehearse all the streets of London with equal importance and frequency, in order to prepare for the license test.. In turn, the more homogeneous distribution of memories creates higher demands for representational space, which might be met with increased synapses or neurogenesis and might lead to increased hippocampal volume.

These results are also compatible with other forms of experience-induced changes in hippocampus size, such as increased hippocampal volume with higher education [41,42]. A large part of educational curricula consists, after all, not only of memorizing a large number of facts but also of re-using them consistently across grades: for example, the chain rule in calculus might be studied in high school and then revisited multiple times in different college classes. Both the large number of notions and their rehearsal contributed to make the distribution of memory probabilities much more uniform (and, therefore, more entropic: Figure 1).

Finally, these results are compatible with the mechanisms proposed by Smith and colleagues [26] for the negative effects of PTSD on hippocampal volume. Because of their intrusive nature, traumatic memories fall naturally in the category of high-frequency memories, which can be encoded in a much sparser way, potentially leading to a reduced number of neurons and synapses maintained in the hippocampus.

Taken together, these results confirm the role of memory frequency in determining the size of a representation and outline a mechanism for how different numbers of neurons can be allocated to different memories. None of these results, however, provide any evidence that these effects are directly linked to the addition to a L1 penalty within the hippocampus, which was introduced specifically to enforce sparseness in memory representations. To test the role played by this mechanism, we will examine how the model simulates a specific clinical population: Autism Spectrum Disorder.

### Simulation 3: Effects of Reduced Lateral Inhibition in ASD

In our autoencoder model, sparseness is achieved by adding a penalty to the loss function. In a real, biological network, however, sparseness would likely be achieved through lateral inhibition, that is, the presence of inhibitory synapses between neurons belonging to the same region. Inhibitory collaterals are a common feature of neuron circuitry, and might be dysfunctional in Autism Spectrum Disorder (ASD). An often-cited post-mortem analysis of individuals with ASD has found a marked reduction in neuronal dendrites neurons in area CA1 of the hippocampus [43]. In addition, inhibitory synapses typically express GABA receptors, and abnormalities in GABA signaling have been often reported in animal models of Autism Spectrum Disorder [44], while magnetic resonance spectroscopy studies in vivo have consistently reported reduced GABA signaling in participants with ASD, especially in the temporal lobe [45]. Similarly, CNTNAP2 genetic variants have been found to represent ASD genetic risk factors in humans, and in mice models, knockout of the CNTNA2 gene results in greater spontaneous activity and reduced inhibition in the hippocampus [46]. This reduced inhibition also aligns with the abnormal association between ASD and epilepsy in humans [47,48], Importantly, neuroimaging studies in humans have also reported a significantly *larger* absolute hippocampal volume in ASD compared to controls, both throughout development [49,50] and in adulthood [51]. This volumetric increase also mirrors the finding that individuals with ASD have higher densities of hippocampal neurons, as reported in a *post-mortem* study [52].

Our model suggests a possible way in which these findings could be related: the reduced GABA signaling and reduced lateral inhibition in ASD would translate to a reduced L1 penalty and, as a consequence, in an increase in the representational costs for encoding memories. In turn, this would lead to less efficient coding, possibly contributing to the enlarged hippocampus observed in ASD.

To test this hypothesis, a series of simulations were carried out using the original training set (with object frequencies of 1, 10, 100, and 1,000) but with the penalty parameter set to λ=0. Under these settings, the only equivalent of lateral inhibition is the implicit threshold that zeroes the activity of ReLU units in the CA3 layer.

The results of the simulations are shown in the red lines of Figure 7, compared to the results of Simulation 1 in gray. As it can be seen, without the resource cost penalty, the model now uses a disproportionately large amount of neurons and incurs a large L1 penalty. To formally analyze the results of these simulations, we applied linear regressions models to both dependent variables, using object frequency (1, 10, 100, or 1,000) and model (neurotypical, with lateral inhibition, vs. ASD, without lateral inhibition) as the fixed-effect independent variables. In both measures, both the object frequency (L1 Penalty: β = –0.02, *p* < 0.0001; Number of neurons: β = –0.03, *p* < 0.0001) and the model type (L1 Penalty: β = 94.55, *p* < 0.0001; Number of neurons: β = 57.08, *p* < 0.0001) had significant effect, as did as their interaction (L1 Penalty: β = 0.003, *p* = 0.003; Number of neurons: β = 57.08, *p* < 0.001). A follow-up analysis revealed that, in the ASD-like model without lateral inhibition, greater object frequency still leads to significantly smaller penalties and number of neurons (β < –0.01, *p* < 0.05). This finding demonstrates that the autoencoder model would still lead to sparse representations for the most frequent objects even without the additional L1 penalty. This is perhaps not surprising: as noted above, the L1 penalty is only one of the mechanisms that induce representational costs. However, the results also show that without the L1 penalty, the effect of object frequency is greatly reduced and the overall size of hippocampal representations increases dramatically.

**Figure 7:**
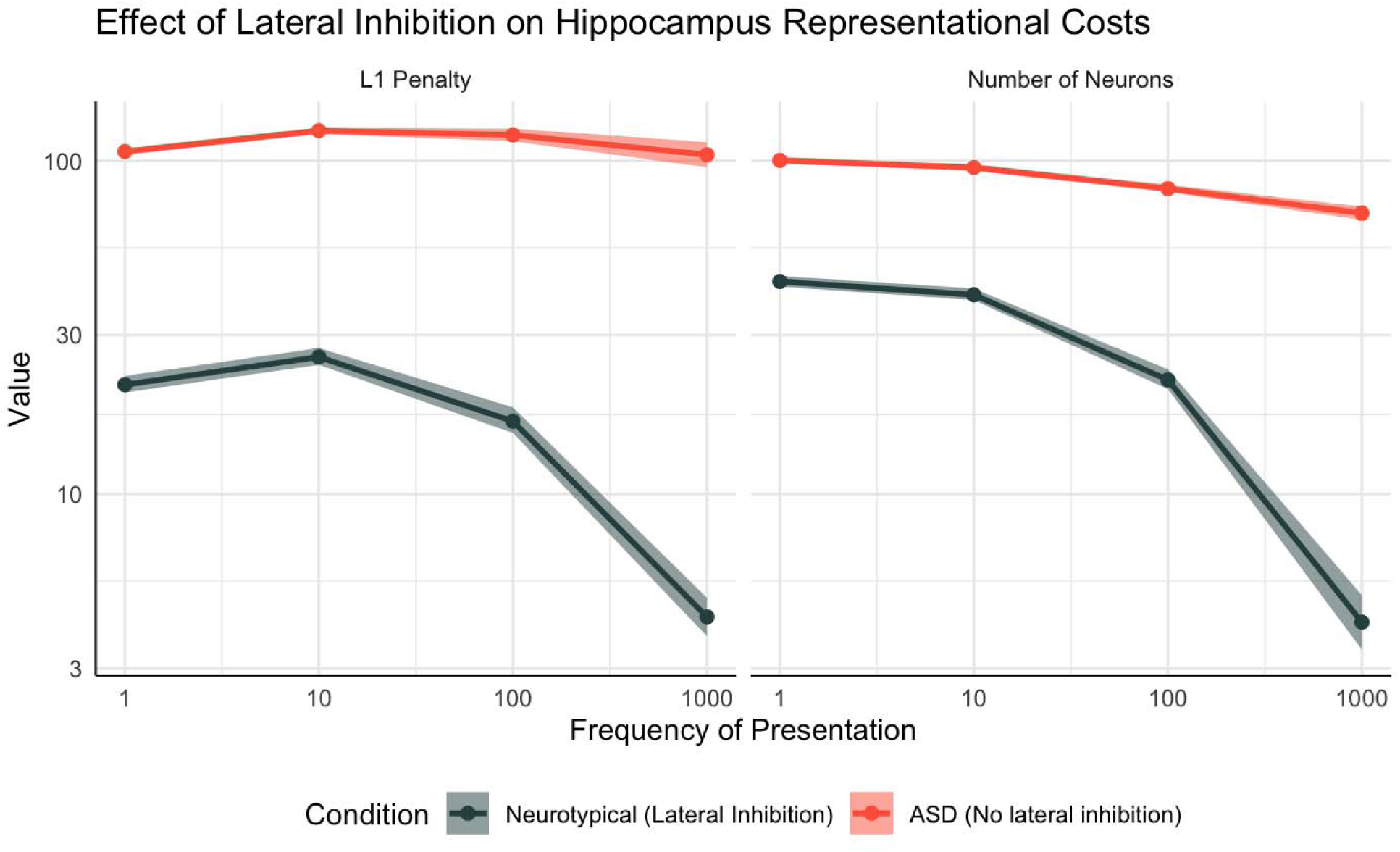
A comparison of the L1 penalty (left) and representational costs (right) in a neurotypical model of the hippocampus (slate color, same as Simulation 1) and a in model where the L1 penalty parameter has been set to zero to simulate impaired lateral inhibition in ASD (red).

#### Neural Activity and Memory Function in ASD

These results suggest that, indeed, if ASD does manifest with reduced inhibition or, at the cellular level, with reduced collaterals, then an enlarged hippocampus size is to be expected, based on the additional representational demands. In addition to the greater hippocampal volume, our model makes an additional prediction: since reduced inhibition is associated with memory representations spread over a larger number of neurons, it should also result in greater neuronal activity when memories are formed or retrieved. Although this prediction has never been tested directly, neuroimaging studies of ASD patients performing memory tasks have reported increased hippocampal activity during memory retrieval [53,54]; recent meta-analysis of multiple neuroimaging studies (including evidence from other imaging modalities, such as EEG and MEG) has also found that ASD is consistently associated with greater medial temporal lobe activation during memory recall, especially in the left hemisphere [55].

Despite the evidence of structural and functional changes in the hippocampus, the clinical evidence is mixed as to whether individuals with ASD exhibit accompanying impairments in long-term memory function or not; for example, a recent meta-analysis [56] of 64 behavioral studies found consistent deficits in ASD for *short*-term memory, but inconsistent and small effects for *long*-term memory. The setup of our simulations is poorly suited to investigate a direct relationship between representation size and recall precision: under different conditions (e.g., greater or smaller L1 penalty), the training algorithm will lead to different object representations but, because of its adaptive nature, will also strive to maintain equal performance. It is possible, however, to indirectly examine the effects of representation on memory performance by artificially damaging the model after learning, and measuring the model’s recall accuracy under different conditions. The next two sets of simulations will address this point by simulating the effects of neuronal damage, such as those seen in dementia, and their interaction with reduced inhibition in ASD.

### Simulation 4: Differential Impairment of Recent vs. Past Memories in Dementia

As noted in the introduction, while changes in hippocampal volume are not consistently associated with changes in long-term memory function, *some* forms of pathological reductions in hippocampal size lead to distinctive deficits in memory. This is the case, for example, of neurodegenerative diseases such as Alzheimer’s Disease (AD) where neuronal loss afflicts long-term memory by harming the engram associated with the memory [57,58].

To simulate the effects of AD, we ran a fourth set of simulations. These simulations are identical in nature to Simulation 1 but with an additional manipulation: After completing the training phase, the model’s hippocampus was artificially damaged by applying a binary mask to the activation of its units. Binary masks were generated by first creating a unary vector of 256 elements and randomly setting a percentage of its elements to 0; the proportion of units set to 0 represents the proportion of neuronal damage in the hippocampus, and was parametrically 0.1 to 0.9. After every simulated lesion, the model’s recall accuracy was tested again, and the correlation between the original and the recalled stimulus was recorded and transformed into a *Z*-score. Figure 8 provides examples of the recall of four of different frequencies after 50% of cells in the CA3 were lesioned (lesions cells are shown in red). The example illustrates cases in which the damage barely affects the object presented only once; significantly affects the recall of the object presented 10 times, and leaves the recall of the other two objects unscathed.

**Figure 8:**
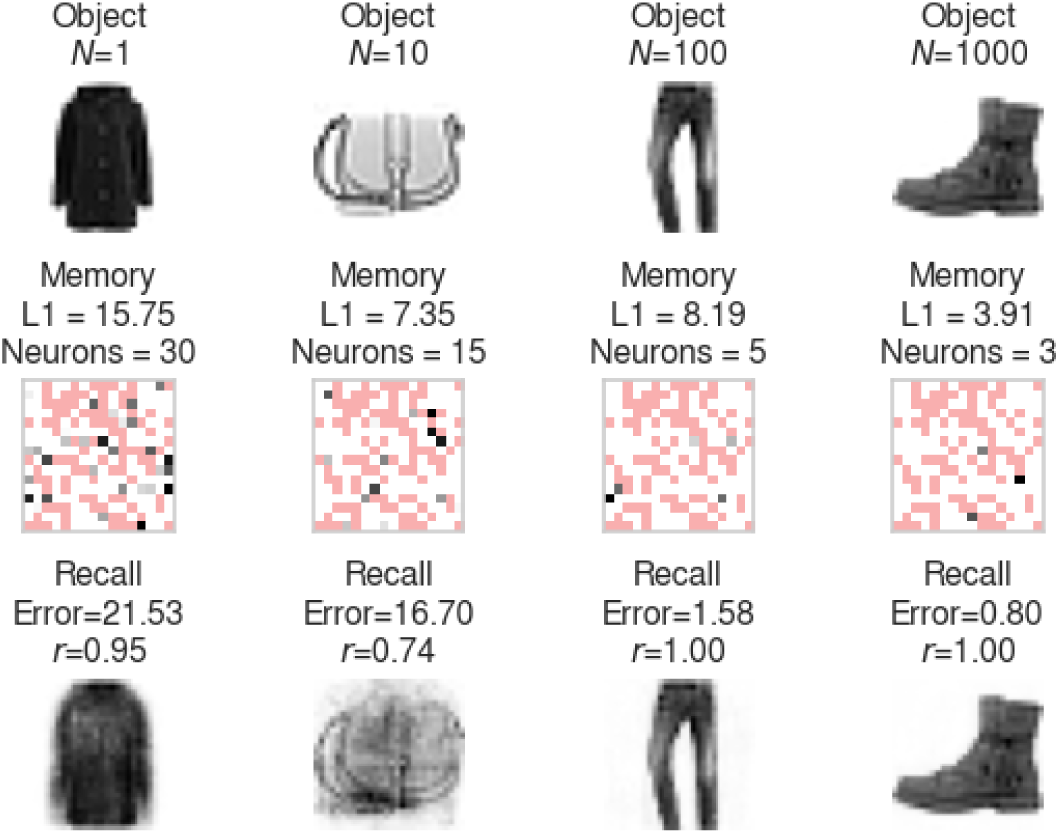
(Top) Four example stimuli that were repeated 1, 10, 100, or 1,000 times in the training set. (Middle) Corresponding CA3 representations of the stimuli; cells damaged to simulate AD are shown in red; (Bottom) Recalled stimuli reconstructed by the decoder from the CA3 representations, together with their recall correlations and the squared pixel error.

Figure 9A provides an overview of the results of the 50 different runs, plotting recall accuracy across different object frequencies for different levels of damage. For the purpose of statistical analysis, the recall correlation values are transformed into equivalent *Z*-score—higher *Z*-scores correspond to higher correlations. Consistent with the observed symptoms of dementia, less frequent memories are more affected, even at the lowest levels of neuronal damage, than the more frequent ones, which often remain comparatively well preserved even at lower levels of neuronal integrity.

**Figure 9:**
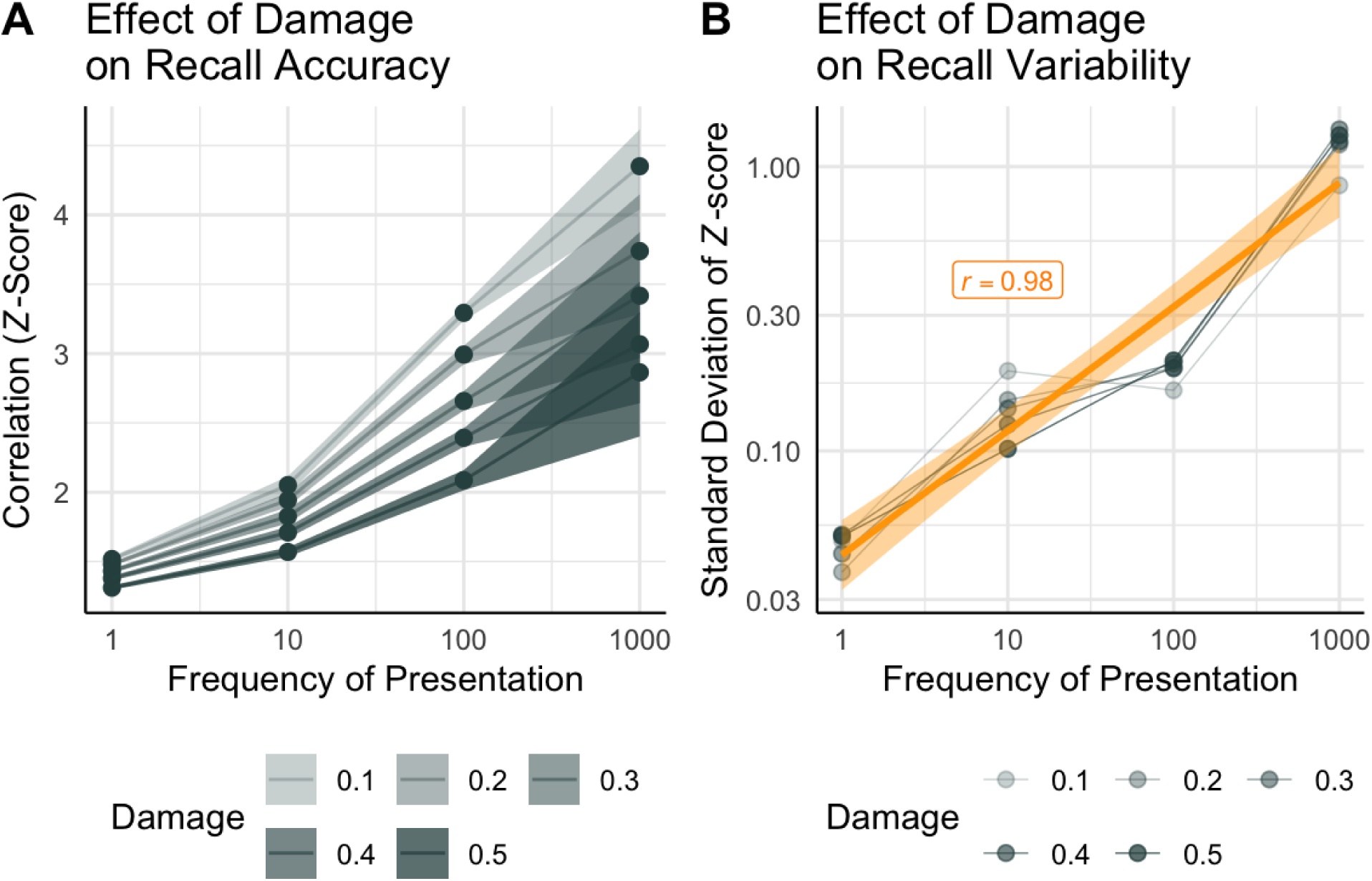
(A) Effects of neural damage on recall accuracy. Lines and ribbons indicate means +/– 99% confidence intervals; levels of transparency indicate different proportions of damaged CA3 cells. (B) Effects of neural damage and stimulus frequency on recall variability, measured as the standard deviation of the Z-scores. Orange line and ribbon represent regression line + SE.

To further analyze these patterns, recall correlations were analyzed with a regression model that included object frequency and proportion of damaged cells as the main factors as well as their interaction. Greater damage negatively impacted recall, decreasing the similarity between the encoded and the recalled stimulus (β = –1.59, *p* < 0.001). Greater frequency resulted in greater recall correlation overall (β = 0.002, *p* < 0.001) and in a negative interaction with damage (β = –0.002, *p* < 0.001), such that the benefits of frequency were reduced when the proportion of damaged cells were higher.

#### Preservation of Older Memories in Dementia

It is well known that, while AD profoundly affects memory, not all memories are equally affected. Older episodic memories are, in general, better remembered [59–61], Being older, these preserved memories are also more likely to be revisited, thus increasing their overall frequency. Thus, the pattern of results in Figure 9A are broadly compatible with the existing literature.

These results, however, do not explain *why* older and more frequent memories are more likely to be spared by neuronal damage. A possible mechanism is that older memories (or pleasant emotional memories, such as beloved songs: [62]) would be revisited more frequently, and thus recruit a smaller number of neurons and synapses; this, in turn—which would make them more likely to be missed by the neuronal damage. The same mechanisms, however, makes older memories more liable *if* they are affected by the damage; an engram containing only three neurons (such as the shoe stimulus in Figure 3) is more likely to be spared but, if one of its neurons is damaged, its recall is also more likely to be severely compromised.

Computationally, this implies that, as frequency increases, recall accuracy should improve but its variability should also increase. Such an effect is visible in the increased ribbon widths of Figure 9A. To analyze this prediction quantitatively, we calculated the standard deviation of the *Z*-scores for each level of damage and object frequency; the results are shown in Figure 9B. As the figure makes it clear, the standard deviation of the recall accuracy varies minimally across proportions of cells damaged, but is greatly affected by object frequency. In fact, standard deviation and frequency are almost perfectly correlated (*r* = 0.98, *p* < 0.001).

### Simulation 5: Reduced Resilience to Neural Damage in ASD

The results of Simulations 3 and 4 suggest that an additional advantage of the efficient sparse coding of memories is to buffer against neuronal death. Thus, a straightforward prediction of our model is that conditions that systematically lead to more expansive and less sparse memory representations, such as ASD, would also make memories more susceptible to neural damage, such as the loss of neurons caused by AD. Consistent with this hypothesis, although ASD *per se* is not associated with either large or systematic changes in long-term memory function [56], dementia has a higher prevalence in individuals with ASD than in neurotypical controls [14,63].

To test this whether this finding can be reproduced by the model, we conducted a second series of lesion simulations, identical to the ones in Simulation 4 but with the model’s λ parameter set to λ = 0, mirroring the settings used to capture the effect of ASD in Simulation 3. Figure 10 provides a summary of the results, with the left panel reproducing the data from Simulation 4 (in gray) and the right panel showing the new results from a ASD-like model (that is, with no L1 penalty; in red) under the same conditions.A side-by-side visual comparison immediately suggests that, compared to the neurotypical model, the ASD model exhibits lower recall accuracy, especially at higher frequencies, for all levels of neuronal damage.

**Figure 10.**
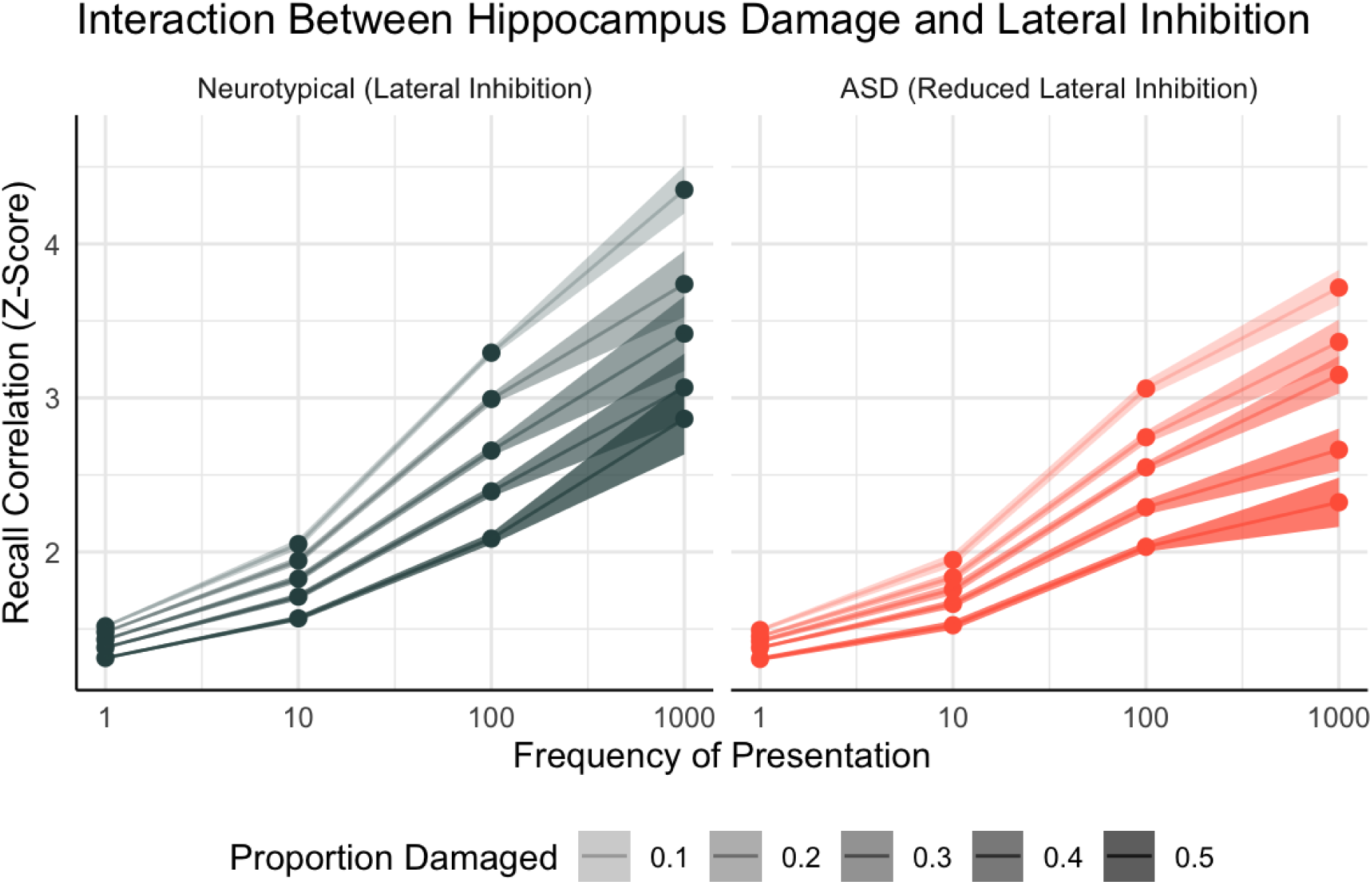
The Z-score recall correlations in a neurotypical model (with L1 penalty, mimicking lateral inhibition in the hippocampus; left) and in a model of Autism Spectrum Disorder (without L1 penalty, mimicking reduced lateral inhibition) across different proportions of damaged hippocampal cells. Line and ribbons indicate means +/– 99% confidence intervals; different degrees of transparency indicate different proportions of damage.

To verify whether the difference was significant, the results were analyzed using a linear regression model whose factors included object frequency, proportion of damage, their interaction, and the type of model (neurotypical vs. ASD). All of these factors were found to be significant in the statistical model. Specifically, greater object frequency was associated with better recall (β = 0.002, p < 0.001); greater damage was associated with worse recall (β = –1.38, p < 0.001); the effect of damage was reduced when objects were encoded more frequently (β = –0.002, *p* < 0.001); and, finally, the neurotypical model was found to have better recall overall (β = 0.17, *p* < 0.001). As predicted, when the model is provided with the L1 penalty, it is consistently less affected by damage across all levels of stimulus frequency.

In Simulation 4, it was hypothesized that the model’s resilience to damage was coming from the “hit or miss” effect of neuronal damage to the extremely sparse representations induced by the combined effect of frequency and the L1 penalty. As noted above, these simulations give us a way to empirically test this prediction. If our reasoning is correct, the distribution of recall accuracies for the neurotypical model should look markedly different from the accuracies for the ASD model. Specifically, we would expect the distributions of recall accuracies to be approximately normal or unimodal in the ASD model, because damage is more likely to impact representations of even the most frequently encountered objects. The sparse representations in the neurotypical model, on the other hand, should be associated with more extreme, bimodal distributions, since neuronal damage is likely to either severely impact the representation or to leave it completely unscathed.

To this point, Figure 11 illustrates the distributions of recall accuracies for the most frequently seen object (frequency = 1,000) in the two models across all levels of neuronal damage. It is clear how, for every level of the damage, the distribution of accuracies for the neurotypical model always has at least two peaks, one very close to zero (corresponding to the case in which the representation has been obliterated by damage) and one to the right of the peak of the ASD model (corresponding to the case in which the representation is either unscathed or minimally affected).

**Figure 11:**
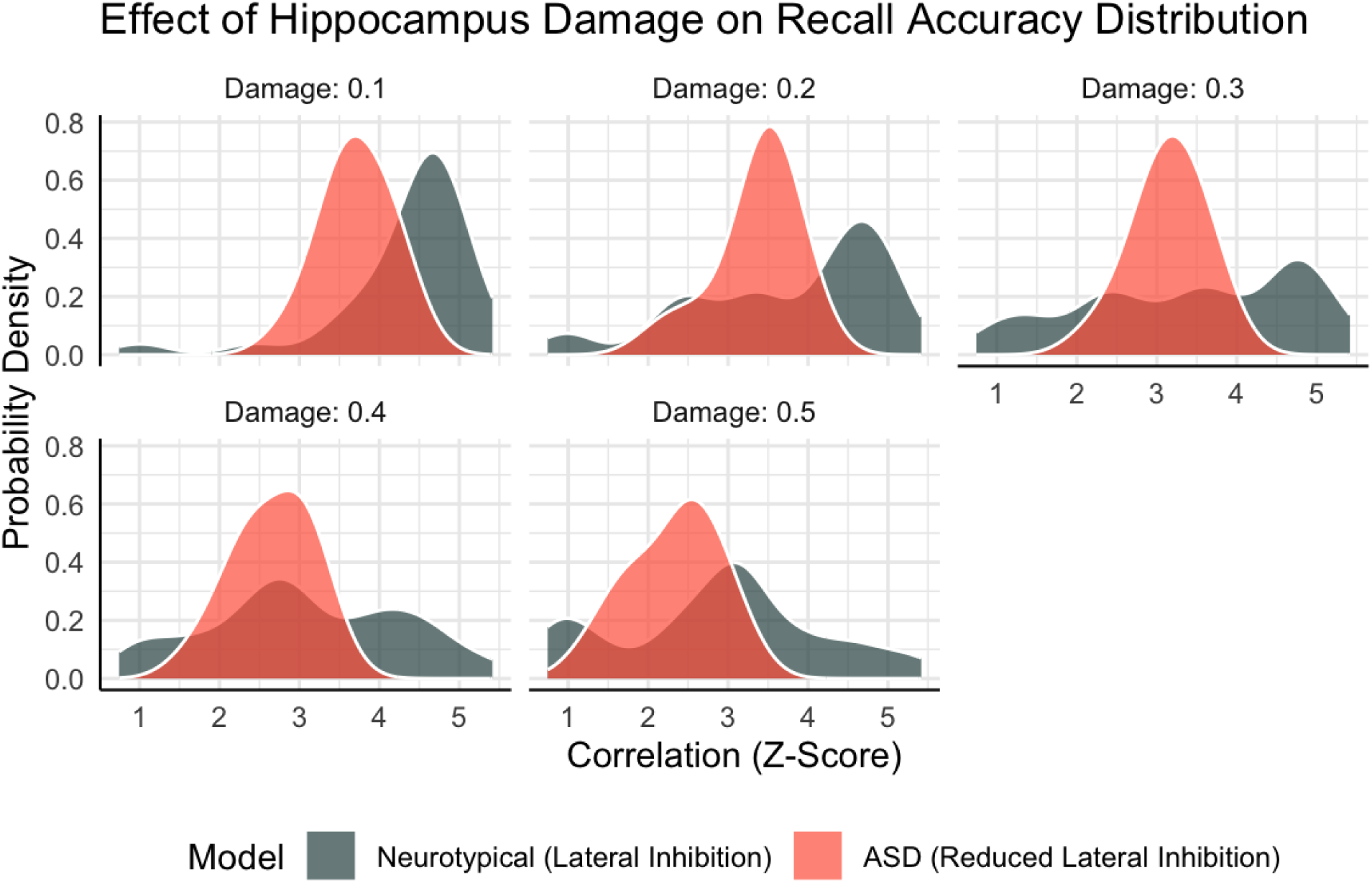
Distribution of accuracies for the most frequently presented object (frequency = 1,000) across different proportions of neuronal damage in the neurotypical (i.e., including the L1 penalty; slate gray) and the ASD (i.e., with no L1 penalty; red) models. Higher z-scores represent higher correlations and, thus, greater recall accuracy.

## Discussion

This paper has proposed a theory that reconciles the puzzling findings in the literature regarding HPC size and memory functions across some clinical and neurotypical populations. Specifically, they can be explained by analyzing the behavior of an autoencoder network model of the hippocampus, where the loss function incorporates a resource cost. The resource cost is implemented as a L1 penalty for hippocampal activation, and induces sparse representations in a way that is adaptive and efficient according to information theory: that is, most frequent stimuli are given sparser representations that involve a smaller number of neurons and synapses. These representations are more in a form that is consistent with the principles of efficient coding and the Free Energy Principle, with most frequently used memories being assigned a sparser code. This provides an objective computational framework for the observed fact that hippocampal size is plastic and adaptable, and responds to the environmental demands. In particular, our findings suggest that the hippocampal size reflects the information content of the stored memories.

Assuming that the hippocampus adapts to environmental demand functions, our model provides an elegant explanation for the observed changes in hippocampal volume when the frequency of memories deviates from uniform, as in the case of PTSD and fewer years of education (Simulations 1 and 2). Abnormally low lateral inhibition or GABA signaling can be interpreted as a reduction in the resource cost penalty, and results in higher representational demands, which could explain the larger absolute volume in the hippocampus in ASD (Simulation 3). Sparser representations are also more resistant with damage, which explains the persistence of older memories in AD (Simulation 4) and the fact that individuals with ASD are more affected by AD (Simulation 5).

Despite being a highly simplified model of the hippocampus, some characteristics of its learned representations are consistent with other findings in the literature. For example, when memories are very frequent, the model tends to form extremely sparse representations. The representation of the shoe object in Figure 2, for example, contains only 3 neurons and, in 14% of our simulations, the hippocampal representations for the most frequent objects was reduced to a single neuron. This extremely sparse representation is reminiscent of the putative “Grandmother cells” that have been anecdotally reported (such as the “Jennifer Aniston neuron”: [64]). Note that our model does not enforce such a “one-hot” form of representation, and most representations are overlapping and distributed. Rather, this representation emerges as a result of environmental pressures to save costs.

One of the consequences of representational sparseness is that most hippocampus neurons would be involved in only a few memory representations. This has been shown, in fact, in a rigorous and systematic examination of place cells in CA3: the authors found that most place cells are preferentially associated with one specific location and that their firing rates across different locations are uncorrelated [36].

Another important facet of our model is that sparseness is not induced by encoding *per se*, but by the interaction between encoding and recall. As noted in the introduction, the hippocampus cannot assign efficient memory codes right away, as they require advanced knowledge of an object’s frequency. In the autoencoder, it is the presence of multiple learning passes and error-driven learning that pushes for sparser and efficient coding over multiple encoding and recall cycles. While this is in sharp constraint to the unsupervised form learning of autoassociator models [65,66], it is also consistent with the larger neural architecture in which the hippocampus is embedded, which forms a continuous loop to and from cortical areas. Similarly, the presence of error-driven learning is compatible with the presence of mesencephalic dopamine projections to the hippocampus [67,68] as well as the fact that dopamine promotes changes in the distribution of phasic activity across hippocampal neurons [69,70].

The fact that the model learns the optimal coding through error-driven learning after recall is also compatible with yet another important neural feature: the spontaneous reply of memories in hippocampal cells. Whether occurring at rest [71,72] or during sleep [73–75], replaying of previous encoded memories is well documented in hippocampal neurons. Recently, Pezzulo et al. [76] have postulated that the functional role of spontaneous memory replay is precisely that of optimizing the internal representation of memories—a function that is entirely consistent with our model.

### Limitations

These contributions notwithstanding, a number of limitations must be acknowledged. First and foremost, the model uses an autoencoder architecture, while hippocampus models are more commonly implemented as autoassociators (e.g., [31,37,66,77–81]). As a consequence, the model is incapable of “one-shot” Hebbian learning and requires error-driven training. he choice of using an autoencoder architecture was motivated by its ability to capture the dynamics of encoding and recall, as well as the interactions between the hippocampus and cortical areas. However, an ideal model should integrate both autoencoder and autoassociator architectures, and incorporate principles of Hebbian learning within the hippocampus[39].

The model is also silent about some fundamental distinctions between different types of memory representations. One of these is the lack of a distinction between semantic and episodic memories, which lies beyond the scope of our model for two reasons. Compared to semantic memories, episodic memories are characterized by their larger spatiotemporal context, which makes them autobiographical in nature. In contrast, our hippocampus model is fed only limited amounts of visual information about specific stimuli, and no general episodic context is provided. Under these circumstances, the model’s memories can be interpreted as semantic (e.g., notions learned at school or streets in London) or episodic (e.g., memories of traumatic events or old memories in AD).

It has been suggested multiple times that, although memories begin as hippocampal engrams, semantic memories are ultimate represented in cortico-cortical synapses, and can be recalled independently of the underlying hippocampal activity [31,79,82–84]. In contrast, in our model the hippocampus remains an obligatory step for memory recall, which aligns more closely with episodic, rather than semantic, information [58,83], and is consistent with the fact that semantic memories, but not episodic ones, are generally spared in Alzheimer’s Disease [60].

A related limitation is that, although we have made attempts to relate the results of our model’s simulations with the effects of traumatic memories, the restricted nature of the simulations did not allow us to *directly* model emotional contributions to memory. The effect that emotion has on memory is believed to be implemented, at the neural level, by the modulatory activity that the amygdala exerts on the hippocampus [85–88]; while these projections can be added to our model, they do represent a significant departure from the original autoencoder architecture, and would require additional assumptions.

Finally, it should be noted that virtually all of these limitations are due to the restricted scope of the model and of its simulations; the model could certainly be expanded in the future to implement additional mechanisms to simulate one-shot, associative learning or to tackle the semantic and emotional memories.

## Materials and Methods

### Model Implementation

The model was implemented in Keras [89] with an underlying TensorFlow [90] engine. In addition to the seven layers depicted in Figure 2, the full model contains 4 additional layers that perform purely technical operations, such as reshaping inputs and outputs and computing penalty terms for the cost functions (see below); although necessary for the model to work, these layers are not relevant for its biological implementation. Altogether, the model has a total of 1,347,282 trainable parameters.

In all simulations, the model was trained using the stochastic gradient descent with adaptive moment estimation algorithm (ADAM: [91]). As noted in the Introduction, the loss function *L* included an *accuracy cost* term (the mean squared difference between the activations of each input neuron and the each corresponding output neuron) and a resource cost (the sum of the activations of each neuron in CA3). Note that the resource cost penalty is equivalent to the L1 penalty used in regularization methods, such as LASSO [92]. This particular penalty term was chosen because, unlike other penalties, it is mathematically proven to force its terms to zero, thus reducing the number of active neurons within a representation.

### Dependent Variables

For each of the 1,111 objects in the training set, three dependent variables were computed. Two variables measured the sparseness of the hippocampus representation: the value of the resource cost L1 penalty and the total number of active neurons (that is, with activation *y*_CA3_ > 0) in the hippocampal CA3 layer. The remaining variable measured the model’s recall accuracy, which was operationalized as the Pearson correlation coefficient between the encoded and recalled image. Because correlation coefficients are bounded and not suitable for statistical analysis, the recall accuracy was transformed into corresponding *Z*-scores using Fisher’s *r*-to-*Z* transform: *Z* = [log(1 + *r*) - log(1 – *r*)] / 2.

## Notes

https://osf.io/wxh2r/

### Competing Interest Statement

The authors have declared no competing interest.

### Summary of Updates

Revised framework, clarified structure, expanded the analysis.

